# Altered dynamics of ATP synthase cause functional irregularities in senescent hiPS-cardiomyocytes

**DOI:** 10.1101/2024.06.04.597306

**Authors:** Silke Morris, Isidora Molina-Riquelme, Gonzalo Barrientos, Frank Schmelter, Stefan Peischard, Guiscard Seebohm, Verónica Eisner, Karin B. Busch

**Affiliations:** Institute of Integrative Cell Biology and Physiology, Schlossplatz 5, Faculty of Biology, University of Muenster, 48149 Muenster, North-Rhine-Westphalia (Germany); School of Biological Sciences, Pontificia Universidad Católica de Chile, Avda. Libertador Bernardo O’Higgins 340 Santiago de Chile (Chile); Institute for Genetics of Heart Diseases (IfGH), Department of Cardiovascular Medicine, University Hospital Münster, D-48149 Münster, North-Rhine-Westphalia (Germany)

**Keywords:** Senescence, human induced pluripotent stem cell derived cardiomyocytes, ATP synthase, function and dynamics, bioenergetic efficiency, cristae ultrastructure, cardiomyocyte contractility

## Abstract

Heart disease is the leading cause of death in the elderly population and the heart is a highly energy-consuming tissue. Aging-related heart failure is often driven by energy depletion in cardiomyocytes (CM), which rely on their abundant, cristae-dense mitochondria for ATP production. ATP synthase, localized along the cristae rims, plays a critical role in energy conversion, but the connection between its organization and function remains unclear. Here, we explored the spatiotemporal organization of ATP synthase in senescent CM at the level of individual complexes. Using single-molecule localization and tracking microscopy, we observed reduced enzyme mobility within the cristae, coinciding with decreased ATP synthase activity, despite a stable resting mitochondrial membrane potential. This reduction in activity was independent of changes in ATP synthase expression or dimerization. Electron tomography revealed an increased prevalence of curved inner membranes and fenestrated cristae in senescent CM, explaining the reduced enzyme mobility. Senescent CM displayed irregular autonomous and paced beating patterns. These abnormalities suggest that impaired cardiac function is directly driven by disrupted energy metabolism, rooted in the suboptimal organization and function of ATP synthase in altered cristae.

## 1. Introduction

Cardiovascular diseases (CVDs) are tightly coupled to aging and the leading cause of death in the elderly population, accounting for almost 40 % of all age-related ailments [1]. The process of aging and the onset of age-related CVDs are strongly linked to reduced energy metabolism and decreased mitochondrial function [2]. Mitochondria are responsible for a multitude of tasks ranging from signal transduction [3] to induction of apoptosis [4]. But first and foremost, they are known as the powerhouses of the cell. In a healthy human heart, more than 90% of ATP is produced by mitochondria in a process called oxidative phosphorylation (OXPHOS) [5]. ATP synthesis is carried out by the electron transport chain (ETC) complexes (Complex I-IV) and ATP synthase (Complex V) located in the cristae sub-compartment of inner mitochondrial membrane (IMM) [6]. The cristae are formed by the multiple folding of the IMM. This folding increases the surface area but also poses a physical barrier, the cristae junctions, to the dynamic proteins diffusing within the membrane [7]. Cristae architecture plays a significant role in organization and optimal function of the OXPHOS complexes [8, 9] and, vice versa, mitochondrial function is reflected in the cristae architecture [10]. This is also dynamic due to complex apparent fusion and fission processes, as more recently revealed [11]. Cardiomyocytes (CM) and other muscle cells with high energy demand contain a large number of mitochondria with a high cristae density [12]. Several studies found that the morphology as well as the cristae architecture is changed in aged CM [13, 14]. In a recent study, we evaluated a model of human induced pluripotent stem cell derived CM (hiPSC-CM) [15] and primary rat neonatal ventricular CM (NVCM), in which we induced senescence using mild doxorubicin treatment [15]. Both CM senescence models displayed a compromised IMM ultrastructure with reduced cristae density, as well as altered cristae morphology. We witnessed a higher number of apparently truncated cristae and partially cristae-depleted mitochondria. Here, we investigate the implications of this altered cristae structure on functional ATP synthase organization.

As we recently showed in another cell model, the spatiotemporal organization of the ATP synthase in cristae is tightly coupled to its function [16]. Besides its role as ATP generator of the cell, the ATP synthase can work in reverse, contributes to the mitochondrial permeabilization (permeability transition pore or mPTP) [17], and serves a structural function. Dimers and oligomers of the complex at the cristae tips contribute to the stabilization of the cristae structure [18], where the local pH is determined by ATP synthase forward or reverse activity [19]. Reverse activity of ATP synthase (ATPase activity) is initiated when ETC activity is compromised. Thus, ATPase ensures the stability of the electrochemical membrane potential (Δψ_M_) under hypoxic or glycolytic conditions, when ATP production by OXPHOS is not favorable [20, 21] to guarantee the transport of positively charged proteins and metal ions into the mitochondrial matrix [22, 23]. However, in a previous work, we showed that increased ATPase function is associated with a higher mobility of ATP synthase, suggesting that protein dynamics of ATP synthase is coupled to its function [16]. This was associated with a decrease in the relative amount of dimers. A recent study proposed that defective dimerization of ATP synthase might be the cause for mitochondrial energy deficiency in aged CM in mouse heart [13]. Last, it has to be mentioned that ATP synthase dimers and oligomers have a cristae-shaping function [24]. It is not clarified, whether and how onset of ATPase and increased monomerization is connected to changes in cristae architecture.

Here, we provide a deeper analysis of the spatiotemporal organization of ATP synthase during senescence. Our experimental approach is built upon the results of our previous work, where we observed ultrastructural changes in combination with reduced mitochondrial function and increased ROS production under senescent conditions. We found decreased ATP production in senescent CM and examined Δψ_M_ as well as ETC complex composition and activity as potential causes. Unlike previously reported for aged mouse CM, we found no alteration of the oligomeric states of ATP synthase in senescent human CM. However, mobility analysis of ATP synthase by single molecule tracking revealed a decreased diffusion coefficient of ATP synthase. In parallel, we observed a change in mitochondrial ultrastructure towards increased fenestration of cristae. However, no change in ATP synthase protein levels was observed, leading us to conclude that the altered spatiotemporal organization of ATP synthase in conjunction with the observed change in cristae architecture results in suboptimal ATP production. The resulting energy deficit directly affects the electrical activity of cardiomyocytes.

## 2. Experimental Procedures

### Cell Culture and Differentiation of hiPSC

All cells were routinely cultured in a humidified atmosphere of 5% CO_2_ and 37°C. Senday Foreskin 1 (SFS.1) cells were cultured in FTDA Medium [25] on plates coated with Matrigel (Corning, #354263) with daily medium exchange. Cells were passaged when they reached 100% confluency under presence of Rock-inhibitor Y-27632 (R&D Systems, #1254/10). Differentiation into cardiomyocytes (hiPSC-CM) was carried out according to Peischard *et al*. ([26]. Briefly, cells were seeded at high density of 500 000 cells per well of a 24-well plate on day 0 of differentiation in differentiation medium [27]. Concentrations of BMP-4 were batch dependent and were determined in a first test differentiation. Medium was exchanged daily with fresh TS-ASC medium [27] for 7-10 days until autonomous beating was observed. To ensure differentiation into cardiomyocytes, the WNT pathway was inhibited by adding C59 on days 2 and 3 of differentiation. After differentiation, cells were frozen in liquid nitrogen. For experiments, hiPSC-CMs were thawed and seeded out on plastic coated with 0.2 % gelatin (Sigma Aldrich, #G1393) or glass coated with 0.2 % gelatin and FBS (PAN Biotech, #P30-3031) and cultured for at least 2 weeks prior to experiments.

For ventricular differentiation, beating cardiomyocytes were transferred after 8 days to Matrigel®-coated plates with a layer of gelatine. Cells were then cultured in KO-THAI medium (KO-DMEM with PSG, ITS, 0.2 % HSA, 250 µM phospho-ascorbate, 0.008 % thioglycerol, for an additional 28 days to develop a more mature ventricular phenotype.

### Cardiac Myocyte Isolation and Culture

Cardiomyocytes were isolated from the hearts of neonatal Sprague–Dawley rats as described previously [31]. Rats were bred in the Bioterio of the Pontificia Universidad Católica de Chile. All studies followed the Guide for the Care and Use of Laboratory Animals published by the US National Institutes of Health (NIH Publication No. 85-23, revised 1996) and were approved by Institutional Ethics Review Committee. Briefly, ventricles were dissected, pooled, and myocytes dissociated in a solution of collagenase (Worthington #LS004176) and pancreatin (Sigma Aldrich #8049-47-6). After enzymatic dissociation, the cells were selectively enriched for cardiac myocytes (NVCMs) by being plated in F-10 Ham medium (Gibco #11550-043) supplemented with 10 % (v/v) horse serum (Gibco #16050-114), 5 % (v/v) fetal bovine serum (Gibco #16000-044), penicillin and streptomycin (100 units/ml; Thermo Scientific #15140122). The NVCMs were plated on top of 2 % (w/v) gelatin-coated glass coverslips. Finally, the cells were kept at 37°C in a humid atmosphere of 5 % CO_2_. 5-bromo-2-deoxyuridine (Sigma Aldrich #59-14-3) was added to prevent the proliferation of remaining fibroblasts.

### Induction of Senescence

Senescence was induced by treatment with 50 nM doxorubicin (Doxo, Sigma Aldrich, #D1515) for 96 h according to Morris *et al*. [28].

### Generation of a Stable Cell Line

For generation of an SFS.1 cell line stably expressing the subunit gamma (SUγ) of the ATP synthase fused to the HaLo tag (SUγ-HaLo), a triple vector system was used. The main vector was a well-established stem-cell Tet-On transfection vector, KAO717-pPB-hCMV1-IRES-Venus (A) [29-31], in which we inserted the gene sequence for SUγ-HaLo [16, 32]. We did this by amplifying the sequence for SUγ-HaLo from the original vector using primers with overhanging ends containing restriction sites for *XhoI* and *NotI* (highlighted in bold), which allowed cloning of the sequence into KAO717-pPB-hCMV1-IRES-Venus (B) via the same restriction sites.

Fwd: 5’- **CTCGAG**ATGTTCTCTCGCGCGGGTGTCG -3’

Rev: 5’- **GCGGCCGC** ACCGGAAATCTCCAGAGTAGAC -3’

Expression of SUγ-HaLo was hence controlled by the CMV_min_ promotor. The Venus marker that was encoded on the transfection vector allowed for easy detection of positive clones and was controlled by an IRES2 sequence. The second vector was KA0637-pPgCAG-rtTAM2-IN. This vector contained a G418 resistance gene for selection of positive clones, as well as the rTA transactivator sequence that allows doxycycline-inducible expression of SUγ-HaLo. The last vector of the trio was PB200PA-1 (C). This encoded the PiggyBac transposase that enabled stable insertion into the genome of transfected hiPS cells.

We transfected SFS.1 cells with Lipofectamine 3000 (Thermo Fisher Scientific, #L3000-001) according to the manufacturer’s instructions using the three plasmids at a ratio of A:B:C = 15:3:1. Medium was changed after 8 hours. The optimal concentration of G418 was previously determined by generation of a kill curve. We started selection with 50 µg ml^-1^ G418 (Sigma Aldrich, #A1720) 24 h after transfection, then increased this to 150 µg ml^-1^ after 48 h and finally to 200 µg ml^-1^ after 120 h. Selection was considered complete when no more cells were viable in the non-transfected control, which was 12 days post-transfection. Cells were then seeded in a dilution series onto a 96-well plate, which led to monoclonal cultures in some wells. Selection pressure was reduced to 20 µg ml^-1^ for another 5 days before G418 was not added to the culture medium anymore. Monoclonal cultures were expanded and expression was induced with 0.25 µg ml^-1^ doxycycline. Heterogenous expression of Venus (green) and SUγ-HaLo (labelled with HTL-TMR) was confirmed via confocal microscopy. Clones that did not express the two constructs were discarded. Subsequently, a test differentiation into cardiomyocytes was carried out as described above.

Before experiments, expression of SUγ-HaLo was induced by adding 0.25 µg ml^-1^ doxycycline for 96 h.

### Quantitative PCR

Total RNA was extracted from differentiated hiPSC-CMs using the Monarch Total RNA Miniprep Kit (NEB #T2010) and equal amounts of RNA per sample were transcribed into cDNA using the GoScript™ Reverse Transcription Kit (Promega #A5000). Quantitative PCR (qPCR) was carried out on a StepOnePlus™ Real Time PCR System (Thermo Scientific) using the PowerUP™ SYBR™ Green Master Mix (Applied Biosystems #A25741), between 60 and 80 ng of cDNA and 0.1 nM of each forward and reverse primer per reaction. The PCR program consisted of an initial denaturation step at 95°C for 10 min followed by 40 cycles of 95°C for 15 s and 60°C for 60 s. A melting curve was generated after the run (60°C to 95°C at 0.8°C increase per minute). GAPDH served as endogenous control. Primers were synthesized by Eurofins Genomics (Table 1).

**Table 1:**
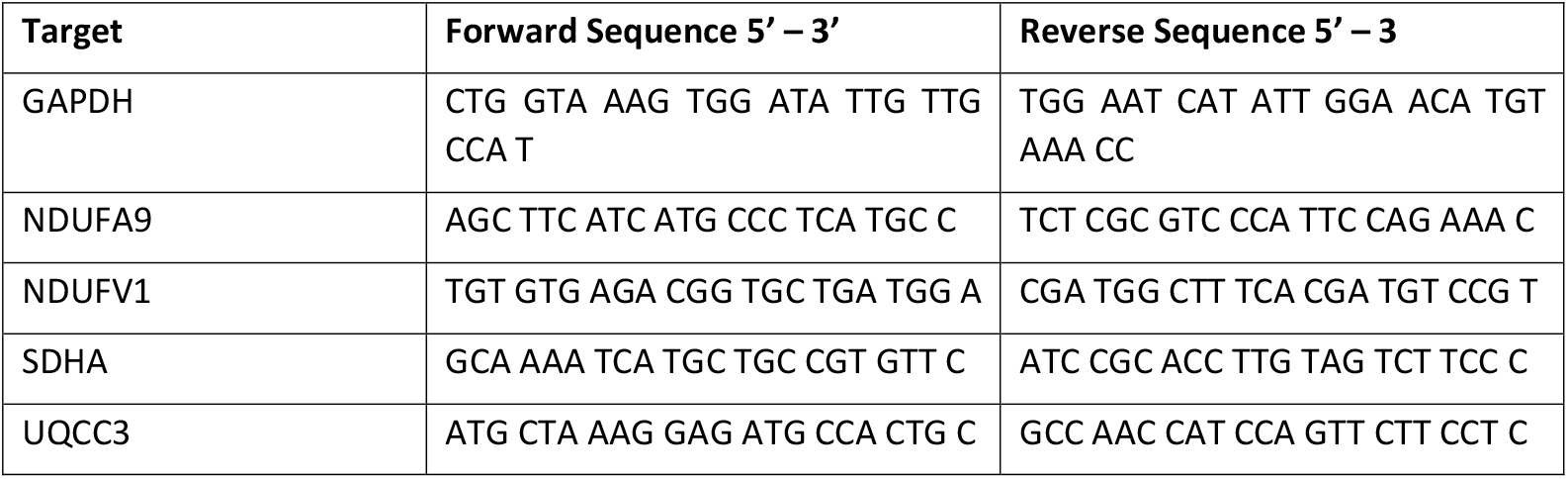
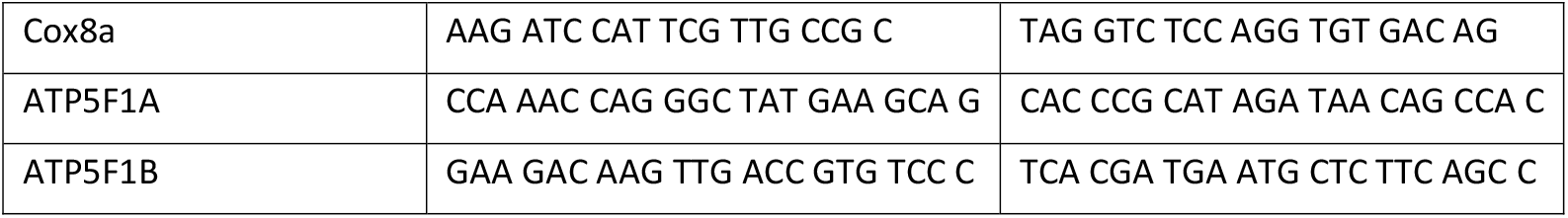
Primers used for qRT-PCR.

### Western Blotting

Cells were lysed in RIPA Buffer (Thermo Scientific, #89901) with added protease inhibitor (Thermo Scientific, #78430). Protein quantification in lysates was performed by modified Bradford assay (Roti Nanoquant, Roth, #K880). Equal protein amounts were supplemented with 4x SDS-PAGE sample buffer and subsequently loaded on a 10 % (w/v) polyacrylamide gel. After electrophoresis, gels were transferred onto a PVDF membrane. Membranes were blocked with 10% milk and immunoblotting was performed. Primary antibody was the OXPHOS rodent WB antibody cocktail (abcam, #ab110413), secondary antibody the goat anti-mouse secondary antibody (Jackson ImmunoResearch, #115-035-068). Visualization of bands was enabled by chemiluminescence (SuperSignalTM West PICO PLUS, Thermo Scientific, #34580) and membranes were stained with Coomassie blue for signal normalization.

### Mitochondria Isolation and Blue Native Gel Electrophoresis

Blue Native gel electrophoresis (BN-PAGE) was conducted according to Wittig *et al*. [33]. SUγ-HaLo SFS.1 and neonatal rat ventricular cardiomyocytes (NVCM) were grown in a T75 culture flask to confluency. After harvesting, the cell pellet was resuspended in ice cold mitochondrial isolation buffer consisting of 300 mM sucrose, 5 mM TES, and 200 µM ethylene glycoltetraacetic acid (EGTA), pH 7.2. Cells were then lysed mechanically by slowly passing the cell suspension six times through a cell homogenizer (Isobiotec) using a 6 µm clearance tungsten carbide ball and glass syringes (Hamilton). The obtained lysate was subjected to centrifugation at 4°C, 800 x *g* for 10 min 3 times to remove nuclei and cell debris. The mitochondria-containing supernatant was subsequently centrifuged at 4°C, 9000 x *g* for 15 min. The resulting pellet contained mitochondria and was resuspended in mitochondrial isolation buffer. Protein concentration was quantified by Bradford assay (ROTI Nanoquant, Roth, #K880.3). 50 µg protein was subjected to another round of centrifugation at 4°C, 9000 x *g* for 10 min and the resulting pellet was resuspended in blue native sample buffer [33] supplied with 4 % digitonin (Sigma Aldrich, #D5628). Samples were separated on a 3 – 12 % polyacrylamide gradient gel (Serva, #43250) alongside a molecular weight marker (NativeMark, Thermo Fisher Scientific, #LC0725). After electrophoresis, the gel was transferred to a PVDF membrane and western blotting was performed using an anti-Halo primary antibody (Promega, #G9211) and a goat anti-mouse secondary antibody (Jackson ImmunoResearch, #115-035-068). For visualization of Complex V, anti-ATPB (subunit β of the ATP synthase, abcam, cat# ab170947) was used as a primary antibody and goat anti-rabbit (Jackson ImmunoResearch, cat# 111-035-045) as secondary antibody. Bands were visualized by chemiluminescence (Super Signal West PICO PLUS, Thermo Fisher Scientific, #34580).

### Seahorse Extracellular Flux Analysis

Seahorse Extracellular Flux Analysis allows the measurement of the oxygen consumption rate (OCR) and the extracellular acidification rate (ECAR) as well as the Proton Efflux Rate (PER) which is derived from the ECAR, while injecting the samples with different reagents, such as inhibitors or substrates of complexes of the electron transport chain. We used the Seahorse Extracellular Flux Analyzer (Agilent Technologies) to perform an ATP rate assay and a complex activity assay.

Differentiated hiPSC-CMs were grown on gelatin-coated 96-well Seahorse plates for at least two weeks before the experiment. On the day prior to the experiment, the Seahorse XF extracellular flux assay sensor cartridge was hydrated with calibrant solution (Agilent Technologies, #100840-000) and incubated at 37°C overnight.

The ATP rate assay was performed according to the manufacturer’s instructions (Agilent Technologies, #103592-100). Briefly, cells were washed with PBS and medium exchanged for Seahorse XF Base Medium Minimal DMEM (Agilent Technologies, #102353) supplied with 10 mM glucose, 1 mM pyruvate, and 2 mM glutamine. The inhibitors were loaded into the ports of the cartridge to obtain the following final concentrations: Oligomycin, 1.5 µM (port A) and Rotenone/Antimycin A, 0.5 µM (port B). The ATP production rate was then determined from the following equations:

glycoATP Production Rate [pmol ATP min^-1^] = glycoPER [pmol H^+^ min^-1^] glycoPER [pmol H^+^ min^-1^] = PER [pmol H^+^ min^-1^] – mitoPER [pmol H^+^ min^-1^]

PER [pmol H^+^ min^-1^] = ECAR [mpH min^-1^] × Buffer Factor [mmol H^+^ L^-1^pH^-1^] × Volume [µL] × K_Vol_ mitoPER [pmol H^+^ min^-1^] = mitoOCR [pmol O_2_ min^-1^] × CCF [pmol H^+^ pmol^-1^ O_2_]

mitoOCR [pmol O_2_ min^-1^] = OCR_basal_ [pmol O_2_ min^-1^] – OCR_Rot/AA_ [pmol O_2_ min^-1^] OCR_ATP_ [pmol O_2_ min^-1^] = OCR_basal_ [pmol O_2_ min^-1^] – OCR_Oligo_ [pmol O_2_ min^-1^]

mitoATP Production Rate [pmol ATP min^-1^] = OCR_ATP_ [pmol O_2_ min^-1^] × 2 [pmol O_2_ pmol^-1^ O_2_] × P/O [pmol ATP pmol^-1^ O]

The complex activity assay was carried out to analyze the activity of individual respiratory chain complexes. This assay took place in the presence of digitonin to allow entry of membrane impermeable compounds. Before the assay, cells were washed with PBS and medium was exchanged for MAS++ buffer. This consisted of 220 mM Mannitol, 70 mM Sucrose, 10 mM KH_2_PO_4_, 2 mM HEPES, 1 mM EGTA, 5 mM MgCl_2_, 10 mM glutamate, 10 mM malate, 0.5 µM FCCP, and 0.00006 % digitonin, pH was adjusted to 7.4 with HCl or KOH. Inhibitors and substrates were the following: Rotenone (CI inhibitor), 0.5 mM (port A), Succinate and glyceraldehyde 3-phosphate (CII substrates), 10 mM (port B), Antimycin A (CIII inhibitor), 0.5 µM (port C), Ascorbate, 10 mM and tetramethyl-p-phenylenediamine (TMPD), 100 µM (CIV substrates, port D). In addition, port D was loaded with 0.2 µg ml^-1^ Hoechst 33342 (Thermo Scientific, #H3570) to allow for determination of the cell count via de Cytation 5 image reader and normalization of the OCR to the cell number. From the resulting graph, data for the CI + CIII/CIV activity can be extracted by subtracting the OCR after CI inhibition with Rotenone from the basal OCR, where substrates for CI have been provided. Similarly, CII + CIII/CIV activity is extracted by subtracting the OCR after CIII inhibition with Antimycin A from the OCR after addition of the CII substrates succinate and glyceraldehyde-3-phosphate. Finally, CIV activity can be determined after addition of the electron donor Tetramethylphenylendiamin (TMPD) that can transfer electrons directly to cytochrome c, which transfers them to CIV. CIV activity is subtracting this value from the CI and CIII inhibited respiration.

### Live Cell Imaging

Fluorescence microscopy was carried out on a confocal laser scanning microscope (Leica TCS SP8 SMD) equipped with a 63× water objective (N.A. 1.2) and a tunable white light laser. HyD’s with GaASP photocathodes were used as detectors. Measurements were performed in a humidified chamber at 37°C and 5 % CO_2_. Membrane potential was determined by staining the cells with 7 nM TMRE (Thermo Scientific, #T669) and 100 nM MitoTracker™ Green (MTG, Thermo Scientific, #M7514) for 30 min and subsequent imaging. Image analysis was carried out in ImageJ by measuring the fluorescence intensity of single cells in both channels. The TMRE signal (excitation: 561 nm, emission: 570-695 nm) was then normalized pixelwise to the MTG signal (excitation: 490 nm, emission: 495-547 nm). In a separate experiment, loss of mitochondrial membrane potential was induced by addition of 10 µM FCCP and time lapses were recorded at an acquisition rate of 1 frame per 3 seconds. Resting membrane potential was then calculated as Δ=I_basal_ - I_CCCP_ using absolute intensity values.

ATP levels were determined by staining with 10 µM of the fluorogenic dye BioTracker ATP-Red (Sigma Aldrich, #SCT045). Before measurements, cells were stained with MTG (100 nM for 30 min). Image analysis was carried out in ImageJ by measuring the fluorescence intensity of single cells in both channels and normalizing the ATP-Red intensity to the MTG intensity.

### Immunofluorescence Staining

For immunofluorescence staining, cells were cultured on cover glasses, washing with PBS was carried out after each step. Prior to staining, cells were fixed with 4 % PFA for 20 min at room temperature. Residual PFA activity was quenched by incubation in 0.1 M Glycine for 10 min. Cover glasses were then incubated in 0.1 % Triton X-100 in PBS for 4 min to allow permeabilization of the cells and subsequently blocked in 5 % normal goat serum in PBS for 30 min. Staining was then carried out by incubating the cover glasses with the primary antibody anti-α-actinin (Sigma Aldrich #A7811) diluted 1:400 in blocking buffer for 60 min and with the secondary antibody pAb to RbIgG, tagged with Alexa Fluor 555 (abcam, #150078) diluted 1:400 for 60 min. Coverslips were then mounted onto microscope slides and left to dry overnight before imaging (excitation: 555 nm, emission: 565 nm – 610 nm).

### Calcein/Co^2+^ quenching assay

The green fluorescent dye calcein enters the cell and mitochondria. MitoTracker™DeepRed (MTDR) stains mitochondria. When Co^2+^ is present, it quenches calcein in the cytosol and in mitochondria, if the mPTP is open [34]. Thus, the MRDR/calcein signal reports mPTP opening. Cells are washed in modified HBSS solution containing 10 mM HEPES, 10 µM GlutaMAX and 100 µM NV118. Calcein (2 µM) and Co^2+^ (Cobalt (II) chloride hexahydrate, 1 mM) are added together with 200 nM MTDR for 15 min at 37°C and 5% CO_2_. Then, cells are washed and imaged. Excitation calcein: 495 nm, emission: 505-636 nm; excitation MTDR: 641 nm, emission: 651-790 nm using a cLSM with white light laser excitation and a water objective (1.2 NA).

### Tracking and Localization Microscopy

Tracking and Localization Microscopy (TALM) was carried out as previously described [35]. Differentiated hiPSC-CM were treated with doxorubicin as described or left untreated. 96 h prior to the experiment, expression of SUγ-HaLo was induced using 0.25 µg ml^-1^ doxycycline. One day before the experiment, cells were transferred onto ibidi glass bottom dishes (ibidi, #81156). For imaging, cells were stained with 1 nM Halo Tag Ligand (HTL) tagged with Janelia Fluor 646 (HTL-JF646, Promega, #GA1110) for 30 min at 37°C. Subsequently, cells were washed three times with PBS and phenolred-free medium was added to the cells for imaging. Cells were imaged at room temperature on an Olympus TIRF 4 Line system equipped with a 100x oil immersion TIRF objective (N.A. 1.49) and a diode pumped solid-state laser (excitation 640 nm, 140 mW, emission filter: 600/37 Brighline HC). Imaging was carried out with a highly inclined and laminated optical sheet (HILO) [36], which was achieved by decreasing the illumination angle just below the critical angle for total internal reflection (TIR). 5000 frames were recorded at a frame rate of 58 Hz (17 ms per frame) with a back-illuminated EMCCD camera (Hamamatsu CMOS Model C14440-20UP, pixel size 130 nm).

The labelling density was low enough to allow the identification of single molecules. To confirm mitochondrial localization of the signals, a cumulative projection of all frames was generated. Localization and generation of single molecule trajectories were carried out based on algorithms described by Sergé *et al*. [37]. These were implemented in a custom-made MATLAB code, SLIMfast (provided by C.P. Richter, University of Osnabrück). Briefly, the point spread function (PSF) was experimentally determined by fitting a 2D Gaussian function over the signals. Based on this, single molecules were localized in each frame with an error probability of 10^−7^. Localizations were then connected between frames to form trajectories. Only trajectories were included in the analysis that consisted of at least 5 steps (85 ms) and the search radius was limited to be between 13 and 130 nm (0.2 and 2 pixels).

### CardioExcyte 96 Analysis

Cardiac-like cells derived from hiPSCs were plated on 0.1 % gelatine-coated NSP-96 plates (Nanion Technologies GmbH) at a density of 30,000 cells per well. Cells were seeded KO-THAI and 10 µM Y-27632. After 24 hours, Y-27632 was removed and 50 nM Doxorubicin was added to a subgroup of seeded cells for senescence induction. Spontaneous contractions were observed by day 5. The NSP-96 plates with hiPSC-derived cardiac cells were placed in the CardioExcyte 96 device (Nanion Technologies GmbH) to monitor cell activity (contraction via baseline impedance) for two hours, with measurements taken every 15 minutes for 30 seconds at 37°C and 5 % CO2. The cells spontaneous activity was measured as well as activity under stress at pacing at 3Hz. Data was processed with CardioExcyte Data Control Software (Nanion Technologies GmbH). Peak-to-Peak time of contraction signals were analysed to verify changes in contractile behaviour of non-treated and Doxorubicin-treated cells.

### Statistical Analysis

For statistical analysis, data was tested for normal distribution (Shapiro-Wilk). In case of normal distribution, statistical significance was determined by One-Way-ANOVA followed by Tukey’s Range Test. For non-parametric testing, Kruskal-Wallis ANOVA was used.

## 3. Results

### ATP production is reduced in senescent hiPSC-CM

In our previous study we showed reduced mitochondrial function in senescent hiPSC-CM. Oxygen consumption rate (OCR) measurements showed that basal and maximal respiration were decreased [28]. We asked whether this affects ATP production rates and ATP levels. Therefore, we visualized mitochondrial ATP levels with the fluorescent probe ATP-Red1™ [38] normalized to the MitoTracker™Green (MTG) signal as co-staining. We found a significant reduction of mitochondrial ATP levels from 2.06 a.u. (Median ± 1.68, SD) to 1.63 a.u. (Median ± 1.58, SD, *p<0*.*001*) (Figure 1A). To validate these results, we performed measurements of the OCR as well as proton efflux rate (PER) via extracellular flux analysis (Seahorse/Agilent) in the absence and presence of oligomycin, antimycin A, and rotenone, from which ATP production levels can be derived (Figure 1B). Due to technical issues, it was not possible to normalize this data to the cell number, however, equal distribution of cells was visually confirmed before running the experiment. We saw a decrease of total ATP production from 678 pmol min^-1^ (median ± 172, SD) to 586 pmol min^-1^ (median ± 168, SD, *p<0*.*01*), whereby OXPHOS-related ATP production (MitoATP) decreased from 390 pmol min^-1^ (Median ± 114, SD, *p=0*.*03*) to 368 pmol min^-1^ (Median ± 125, SD), while glycolysis-related ATP production decreased from 281 pmol min^-1^ (Median ± 63, SD) to 241 pmol min^-1^ (Median ± 50, SD, *p<0*.*001*) (Figure 1C). In percentage, the OXPHOS-related ATP production slightly decreased from 56.75 % (Median ± 3.33, SD) to 55.58 % (Median ± 2.13, SD, *p=0*.*03*). This indicates that overall ATP production is reduced in senescent CM and glycolysis could not counteract this energy deficiency. Reasons for impaired OXPHOS-linked ATP synthesis could be a decreased ATP synthesis activity, increased ATPase activity and/or an increased susceptibility of senescent CM to undergo energy collapse by the mPTP [13].

**Figure 1:**
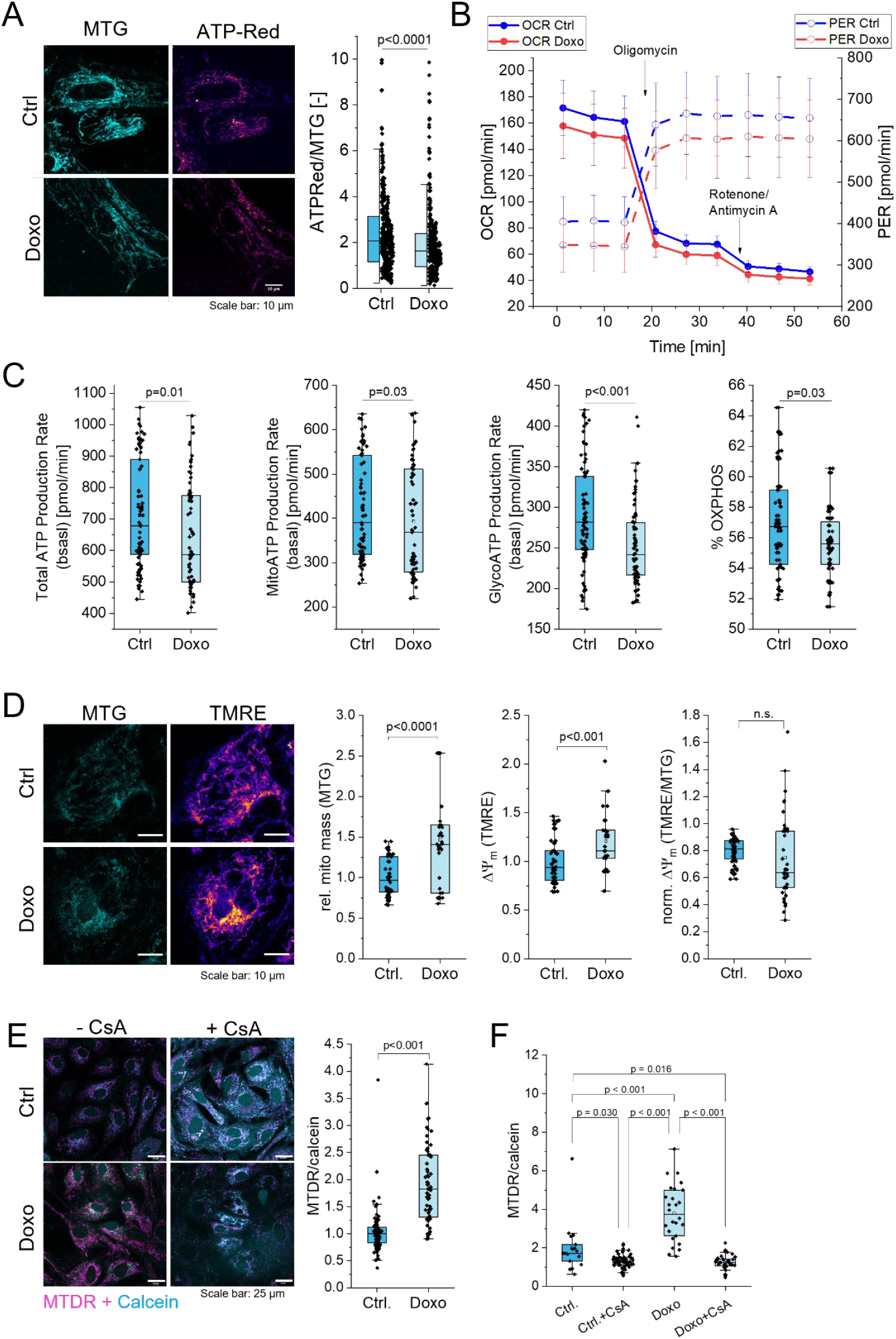
Senescent CM exhibit reduced ATP production and increased mPTP opening. hiPSC-CM were treated with 50 nM doxorubicin for 4 days (Doxo) or left untreated (Ctrl). **(A)** Cells were stained with ATP-Red1™ and MitoTracker™Green (MTG) and fluorescence intensity was measured in both channels. ATP-Red1™ is shown as hot color-coded. The ratio of ATP-Red1™ and MTG intensity was plotted. Statistical test: Mann-Whitney test, n=3 replicates, N=336 single cells. **(B)** Representative time course of ATP Rate Assay carried out via Seahorse extracellular flux analysis. OCR as well as PER were determined in the basal state, after addition of oligomycin and rotenone/antimycin A. **(C)** From **(B)** total, mitochondrial, and glycolytic ATP production rate could be determined, as well as percentage of OXPHOS- and glycolysis-derived ATP production rate. Statistical test: Kruskal-Wallis ANOVA with post-hoc Mann-Whitney. **(A)** n = 3 biological replicates, 468 cells (Ctrl), 483 cells (Doxo). Single points indicate measurements of single cells. n = 3 biological replicates, 77 (ctrl) and 69 (doxo) wells per condition were included in the analysis. MitoATP Production Rate: p = 0.034, %OXPHOS: p = 0.026. **(D)** ΔΨ_m_ was measured by staining with TMRE and MTG for normalization. TMRE staining is shown as hot color-coded. n = 2 biological replicates, 60 cells (Ctrl), 49 cells (Doxo). Single points indicate data of single cells. Statistical test: Mann-Whitney. **(E)** Ratio of Mitotracker™DeepRed (MTDR) and calcein fluorescence in control and senescent CM, and cyclosporin A (CsA) effect on mPTP opening. N=2. 100 cells (Ctrl.), 61 cells (Doxo). Statistic test: Dunn-Sidak. **(F)** CsA inhibits mPTP formation in Ctrl and Doxo cells. Statistic test: Dunn-Sidak. N=1, single points indicate mean values of cells. The horizontal lines in the box blots represent the median values, the error bars denote the standard deviation (SD), the boxes represent the 25^th^ to 75^th^ percentile.

### ΔΨ_m_ is not changed in senescent hiPSC-CM

To test whether reduced mitochondrial ATP synthesis is due to a reduced proton motive force (PMF), we determined ΔΨ_m_ in control and senescent hiPSC-CMs via staining with the membrane potential sensitive dye tetramethylrhodamine ethyl ester perchlorate (TMRE). To normalize on mitochondrial mass, mitochondria were stained with MTG. As before, we found a significantly (1.46 times, *p<0*.*0001*) increased mitochondrial mass [28]. ΔΨ_m_ per cell also increased (Ctrl_Median_ 0.94; Doxo_Median_ 1.11, *p<0*.*001*) (Figure 1D). When the increase in mitochondrial mass was considered, no significant difference in the ΔΨ_m_ per cells was observed, though. We also determined the effect of the uncoupler FCCP on the fluorescence intensity of the TMRE signal. Ctrl. and doxo-treated cells responded with a decrease in Δψ_M_ (Figure S1) demonstrating the specificity of the measurement.

### Senescent CM have a more pronounced mPTP opening

We next asked whether the opening of the mPTP is altered in senescent CM. This was suggested in a recent study, which showed a higher sensitivity for mPTP opening in aged CM [13]. To test for mPTP opening, we applied the calcein/Co^2+^ quenching assay [34]. Cells were exposed to calcein and Co^2+^ and mitochondria were stained with Mitotracker™Deep Red (MTDR). When the IMM gets leaky due to opening of the mPTP, Co^2+^ can enter the matrix and quench calcein fluorescence. The ratio of MTDR and calcein is an indicator for mPTP opening. We observed a significant higher MTDR/calcein ratio in senescent CM due to quenched calcein (Figure 1F). The mPTP antagonist Cyclosporin A (CsA) blocked the mPTP in control and senescent cells confirming the specificity of the assay (Figure 1G). mPTP opening is supposed to reduce the efficiency of mitochondrial ATP production [39].

### Maximal ETC complex activities are not changed in senescent cells

We recently showed that respiration was reduced in senescent hiPS-CM [15]. We wondered, whether this was related to altered maximal ETC complex activity. Therefore, we performed a Complex Activity Assay. The Complex Activity Assay was done with permeabilized hiPS-CM cells and features the injection of inhibitors and substrates of the ETC complexes, allowing the extraction of the OCR related to CI + CIII/CIV, CII + CIII/CIV, as well as CIV alone. It provides maximal activities of the respirasome CI + CIII/CIV, CII + CIII/CIV, as well as CIV alone but this does not necessarily reflect the *in cellulo* situation. An exemplary time course of the experiment is shown in Figure S2. We found no changes in maximal complex activities.

### OXPHOS protein expression is unaltered in senescent cells

Finally, we checked, whether altered expression levels of OXPHOS proteins also contribute to reduced ATP synthesis rates. In a previous study, we showed that respiration was reduced in senescent hiPSC-CM [15]. Here, we asked, whether protein expression levels of OXPHOS complexes were altered. We determined transcript levels by quantitative RT-PCR and protein levels by Western blot. We did not find changes in the transcript levels (Figure S3) nor the protein levels (Figure 2A, B) of representative subunits of the OXPHOS complexes in senescent cells. Overall, we found that the levels of OXPHOS complexes were not significantly altered in senescent cells.

**Figure 2:**
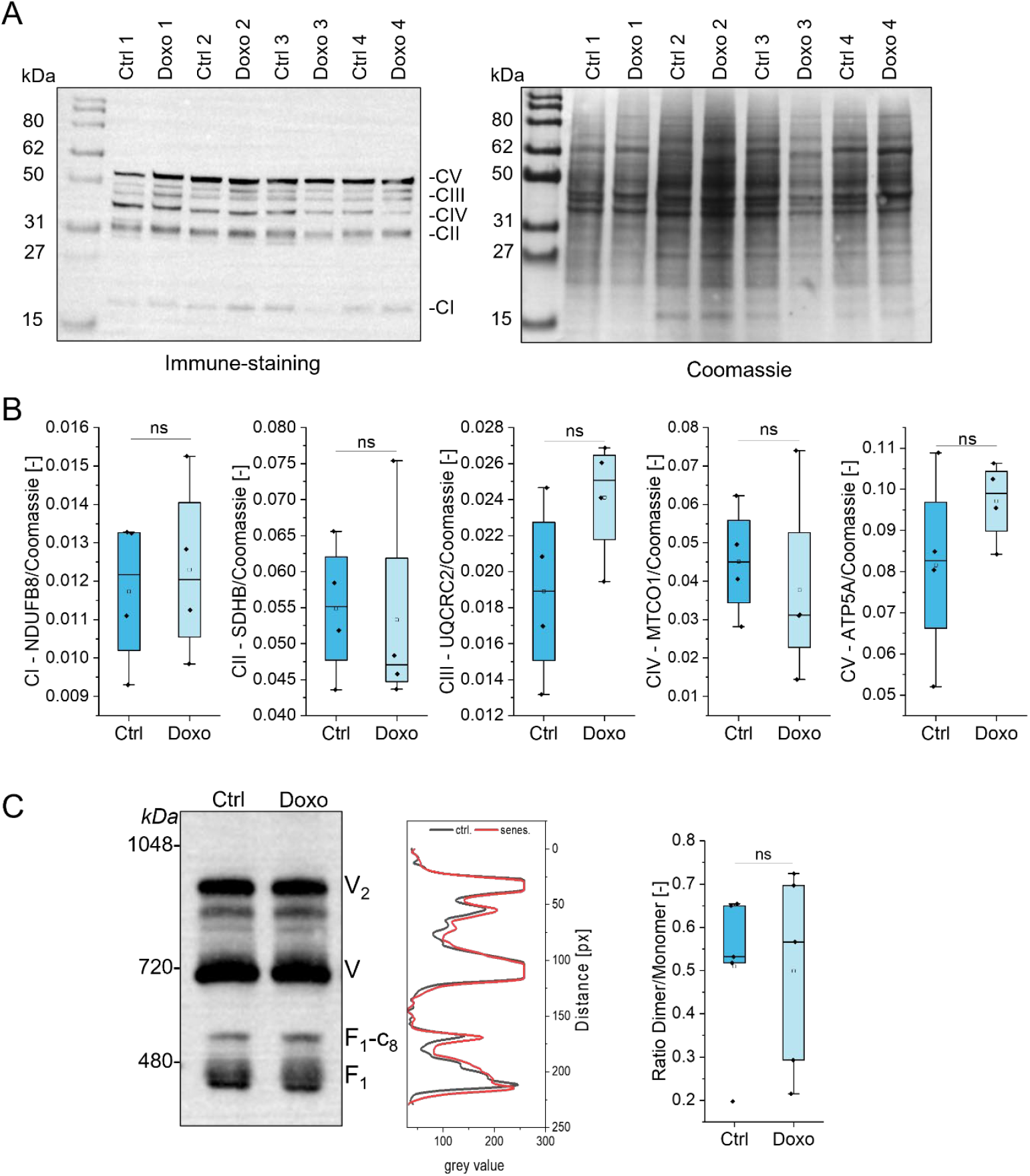
Expression of OXPHOS subunits and ATP synthase organization are not altered in senescent CM. hiPSC-CM were treated with 50 nM doxorubicin for 4 days (Doxo) or left untreated (Ctrl). **(A)** Protein levels of different complex subunits, left: immune-staining (CV: ATP5A, 55 kDa; CIII: UQCRC2, 48 kDa; CIV: MTCO1, 40 kDa, CII: SDHB, 30 kDa; CI: NDUFB8, 20 kDa) right: Coomassie Blue. **(B)** Quantification of **(A)**. Protein levels normalized to total protein load as determined by staining with Coomassie Blue. **(C)** Ctrl and Doxo treated NVCM show no difference in ATP synthase monomer/dimer ratio using an antibody for subunit β. The representative gel shows the dimer (V_2_), monomer (V), as well as F_1_ subunit bound to ring of c-subunits (F_1_- c_8_) and F_1_. Statistical test: Kruskal-Wallis ANOVA because of non-normal distribution of the data. Single points indicate mean values of technical replicates. The horizontal lines in the box plots represent the median values, the error bars denote the standard deviation (SD), the boxes represent the 25^th^ to 75^th^ percentile. Significance: * *p* ≤ 0.05, ** *p* ≤ 0.01, *** *p* ≤ 0.001.

### Levels of ATP synthase and oligomeric organization are not changed in senescent CM

As our immunostaining of subunit α (ATP5A) showed no difference in the protein level (Figure 2B) of ATP synthase, we asked whether the supramolecular organization of ATP synthase was altered in senescent cells and could explain reduced ATP synthesis. The organization of ATP synthase in dimers and oligomers is closely linked to its function [13, 40], whereby dimers and oligomers indicate the ATP synthesis mode [41, 42]. We determined the dimer/monomer ratio by separating ATP synthase complexes via Blue Native Gel-electrophoresis (BN-PAGE) followed by immunostaining (Figure 2C). This was obtained from isolated mitochondria of senescent rat NVCM, since it is not feasible to obtain large quantities of hiPSC-CMs for mitochondrial isolation. The relative amounts of ATP synthase subcomplexes, monomers and dimers were similar between control and senescent CM, other than previously reported, where aging was associated with a loss of dimers [13, 40].

### Senescence reduces the mobility of ATP synthase in cristae

Since the supramolecular organization or total protein levels of ATP synthase were obviously not the reason for the reduced ATP synthase activity, we wondered whether the sub-mitochondrial localization of ATP synthase might be altered in senescent cells. We and others think that optimal coupling and ATP synthesis is related to the specific localization of ATP synthase complexes at bended cristae tips relative to ETC complexes in cristae sheets [42-45]. In order to determine the sub-compartmental localization of ATP synthase in cristae, Tracking and Localization Microscopy (TALM) of single ATP synthase molecules was conducted. This allows to obtain localization and diffusion maps of ATP synthase with nanometer precision [46, 47]. To label ATP synthase, we generated a stable hiPSC line expressing F1-subunit gamma (SUγ) genetically fused to the self-labelling Halo7-Tag. The Halo7-Tag can be fluorescence-labelled via the HaloTag-Ligand [47]. The correct assembly of the tagged SUγ with ATP synthase was tested by immunostaining of a BN-PAGE-blot of isolated mitochondria from undifferentiated WT and SUγ-Halo hiPSC. We found the monomeric form of Complex V (V), with an additional faint band also visible at the size of the dimer (V_2_) (Figure S4A). Staining with an α-Halo antibody confirmed the presence of the labelled subunit at the expected sizes of V and V_2_. In addition, we observed bands for the F_1_ subunit, which consists of subunits α, β, γ, δ and ε, as well as the F_1_-c_8_ subcomplex, which is the F_1_ subunit with a bound ring of c-subunits [48, 49]. We confirmed correct differentiation into CM by visual observation of uniform beating of the cell layer (see Supp Video 1) and by immunofluorescence staining of the sarcomere protein and CM marker α-actinin (Figure S4B). Correct localization of the Halo-tagged SUγ within mitochondria was confirmed by co-localization with TMRE in undifferentiated cells (Pearson mean 0.558 ± 0.086, SD) (Figure S4C).

We used this hiPSC-SUγ-Halo stable cell line to perform single molecule localization and tracking experiments in living CM. The principle of this method is based on the ability to circumvent the diffraction limit [50] by keeping the labelling density very low. This prevents the overlapping of neighboring signals, thus enabling the detection of single molecule signals. Imaging the labelled cells with a high frame rate allows the generation of trajectories from each tagged molecule as long as it is visible in the focal plane. First, signals from individual ATP synthase molecules were fitted by a 2D-Gaussian considering the experimentally determined point spread function. The signal from three individual ATP synthase molecules is exemplarily shown in Figure 3A together with a representation of the localization of a single molecule based on its fitted 2D-Gaussian. We recorded videos with 1000 to 5000 frames. Processing yields the localization of single molecules, which are shown as a cumulative localization map (Figure 3B). The corresponding trajectories are shown in Figure 3C. When an image was taken with 100 ms exposure, the mitochondrial rod-like shape becomes visible (aspect ratio >1) (Figure 3D). The image shows an overlay of the trajectories with the image of mitochondria.

**Figure 3:**
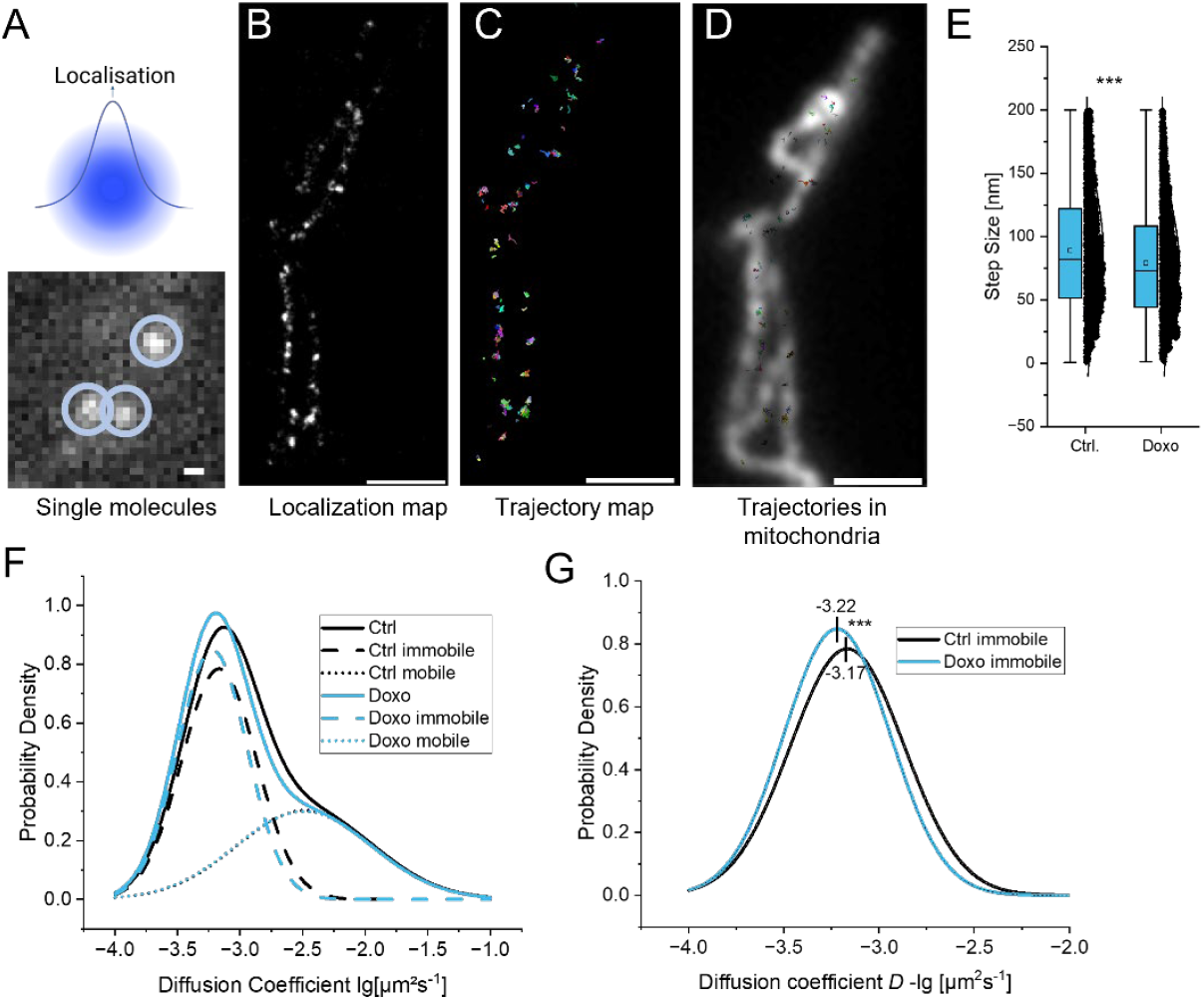
Single molecule localization and tracking of ATP synthase shows decreased mobility of ATP synthase in senescent CM. Stably expressing SUγ-Halo hiPSC-CM were treated with 50 nM doxorubicin for 4 days (Doxo) or left untreated (Ctrl). **(A)** Exemplary single molecules as well as graphical illustration of the point spread function and localization. **(B)** Cumulative image of localizations from 5000 frames. **(C)** Superimposed trajectories from 5000 frames. **(D)** Trajectories overlayed on an image with high exposure time (mitochondria instead of single molecules visible). **(E)** Step size is decreased in doxorubicin treated cells. **(F)** Diffusion coefficients *D* in Ctrl and Doxo cells, shown as probability density distribution of diffusion coefficients of single molecules (solid line). Fitted for two populations, a mobile (dashed line) and an immobile population (dotted line) emerged. **(G)** Probability density distribution of immobile populations fitted with an amplified Gaussian fit. Statistical test: Kruskal-Wallis ANOVA with post-hoc Mann-Whitney because of non-normal distribution of the data. The vertical lines in the box blots represent the median values, the error bars denote the standard deviation (SD), the boxes represent the 25^th^ to 75^th^ percentile. Significance: * p ≤ 0.05, ** p ≤ 0.01, *** p ≤ 0.001. **(A)** Scale bar: 500 nm, **(B-D)** Scale bar: 1 µm **(E)** Scale bar: 250 µm **(F)** n = 3 biological replicates, 7710 trajectories (Ctrl), 2690 trajectories (Doxo). **(E-G)** n = 3 biological replicates, 9220 trajectories (Ctrl), 4576 trajectories (Doxo). Single points indicate measurements of single trajectories.

The step size (Figure 3E) relates to the distance each molecule travels between frames in nm. This is significantly reduced in doxorubicin treated cells from 44.4 nm (Median ± 34.6 nm, SD) to 39.2 nm (Median ± 39.0 nm, SD, *p<0*.*001*). The diffusion coefficient *D*, related to the velocity of single molecules, is displayed as logarithmic values for ease of presentation. The median diffusion coefficient for all molecules decreased from log(−2.99) µm^2^s^-1^ (Median ± log(0.54) µm^2^s^-1^, SD) to log(−3.05) µm^2^s^-1^ (Median ± log(0.54) µm^2^s^-1^, SD, *p<0*.*001*) in senescent cells. This is shown in Figure 3F, which depicts the probability density distribution of the total population (solid lines). This distribution can be fitted for 2 subpopulations, one representing the population of mobile molecules (dotted lines) and one that of quasi-immobile molecules (dashed lines). The mobile populations of both control and doxorubicin-treated cells virtually overlap and show no difference. However, the immobile population of ATP synthase, which is the larger one and indicates dimers and oligomers, is shifted to the left in senescent cells, suggesting a decrease in mobility. This is more clearly depicted in Figure 3G, where we only show the probability density distribution of the immobile population. Summarized, we found a shorter mean step size of ATP synthase in senescent cells and decreased mobility.

### The space of individual cristae is reduced in senescent mitochondria

In our previous study, we observed alterations in cristae architecture in 2D TEM images, suggesting truncation of cristae [15]. This would explain the shorter mean step size and in consequence reduced mobility of ATP synthase. To see changes in cristae architecture in detail, we performed electron tomography of NCVM mitochondria for the control and senescent conditions. An exemplary set of three axial planes for each condition is shown Figure 4. First, we find regular cristae alignment, but also disruptions of crista continuity (Figure 4A, red arrowheads; inset: yellow arrowhead) in the control condition (NVCM). In the senescent mitochondrion, it appears that disruptions were of larger size (Figure 4B, red arrowheads). Such disruptions are only observed in a few planes, because the cristae membrane is continuous above and below them (not shown). This means that these disruptions are structures known as fenestrations, that is, pores in the flat lamellar cristae membrane compartment [51]. Such fenestrations a) reduce the surface area and continuous space of the crista, b) increase the relative contribution of the curved membrane part, and c) shorten the distance between ATP synthase at curved tips and ETC complexes in flat sheets. As the ATP production was lower under these conditions, we conclude that ATP synthesis is not optimal under these conditions. Two possible explanations are presented in the discussion.

**Figure 4:**
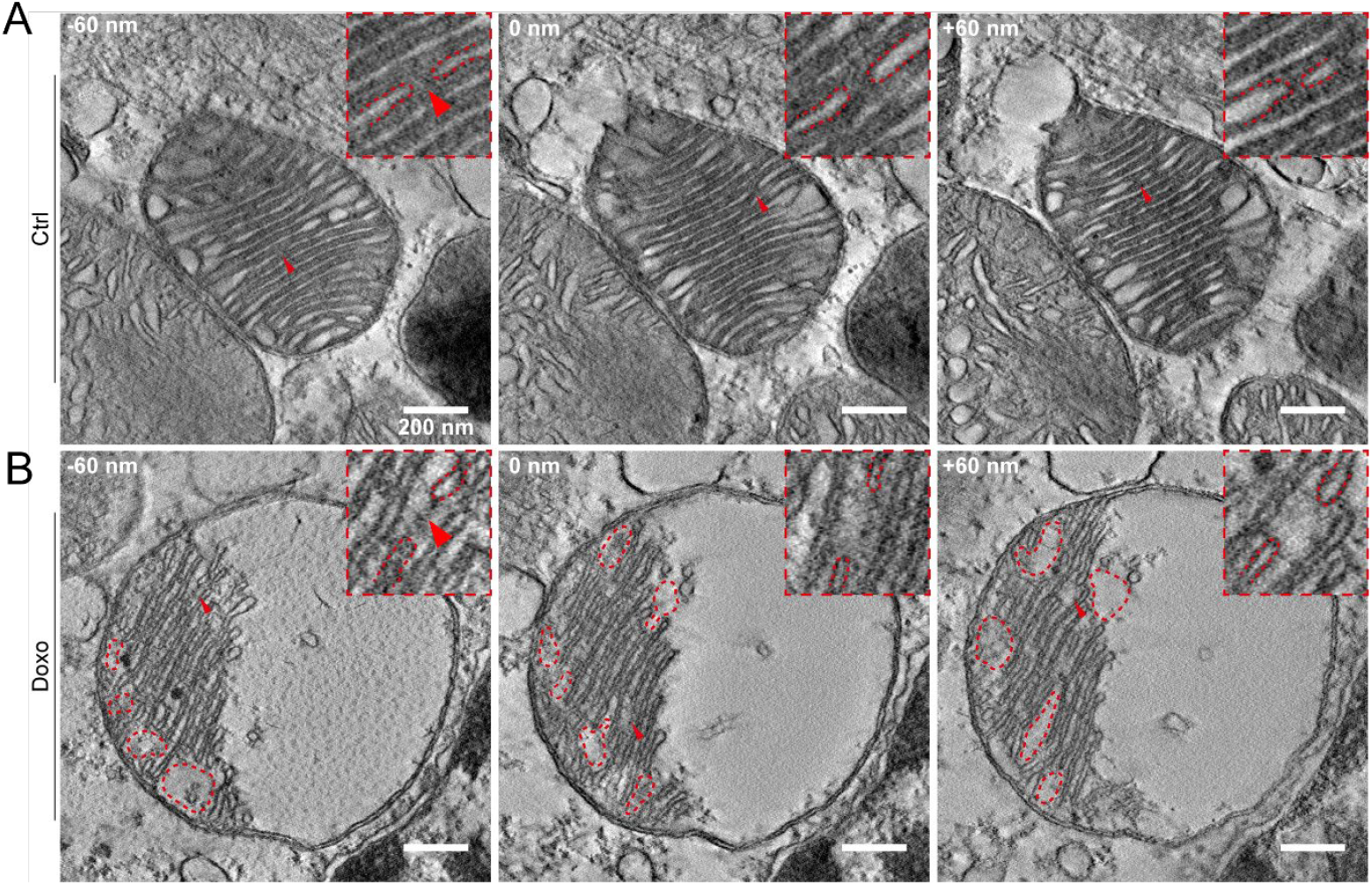
Senescent promotes fenestration of cristae in rat NVCM. **(A)** Mitochondrion from control conditions shown, inset shows detailed view of a crista disruption (red arrowhead). **(B)** Mitochondria under senescent (Doxo) conditions. The inset shows an example of loss of cristae connection, a disruption (red arrowhead). Scale bars: 200 nm. In the figure, there are three axial planes displayed 60 nm apart from one another.

### The contractile activity of senescent cardiomyocytes is more irregular

Finally, we asked how the performance of CM is affected by altered cristae morphology and reduced OXPHOS. We therefore measured the contractility of control and senescent hiPSC-CM with a CardioExcyte 96 device (Nanion Technologies GmbH). Traces of non-treated hiPSC-derived cardiomyocytes were analyzed. The histogram and the plotted Kernel-Smooth-Fit show a clear peak at 1.25 seconds peak-to-peak time (Figure 5A). Analysis of traces of contractility of doxorubicin-treated hiPSC-CM show two peaks at 0.25 seconds and 1.75 seconds peak-to-peak time (Figure 5B). Next, control hiPSC-CM were paced with 3 Hz. The histogram and the plotted Kernel-Smooth-Fit show a clear peak at 1.3 seconds peak-to-peak time (Figure 5C). Furthermore, the fit is slightly shifted to the left compared to non-paced control CM. Finally, doxorubicin-treated hiPS-CM were paced at 3 Hz. Analysis shows two peaks at 0.40 seconds and 1.75 seconds peak-to-peak time (Figure 5D). The appearance of two peak-to-peak times in the senescent cells is a sign of contractile irregularity. The statistical analysis of all conditions shows also significant changes in mean peak-to-peak times. The difference was significant for the contractility of paced control and senescent CM. Paced doxorubicin-treated cells also show a significant difference in beating activity compared to non-paced doxorubicin-treated cells (Figure 5E). In sum, senescent CM displayed a more irregular beating behavior.

**Figure 5:**
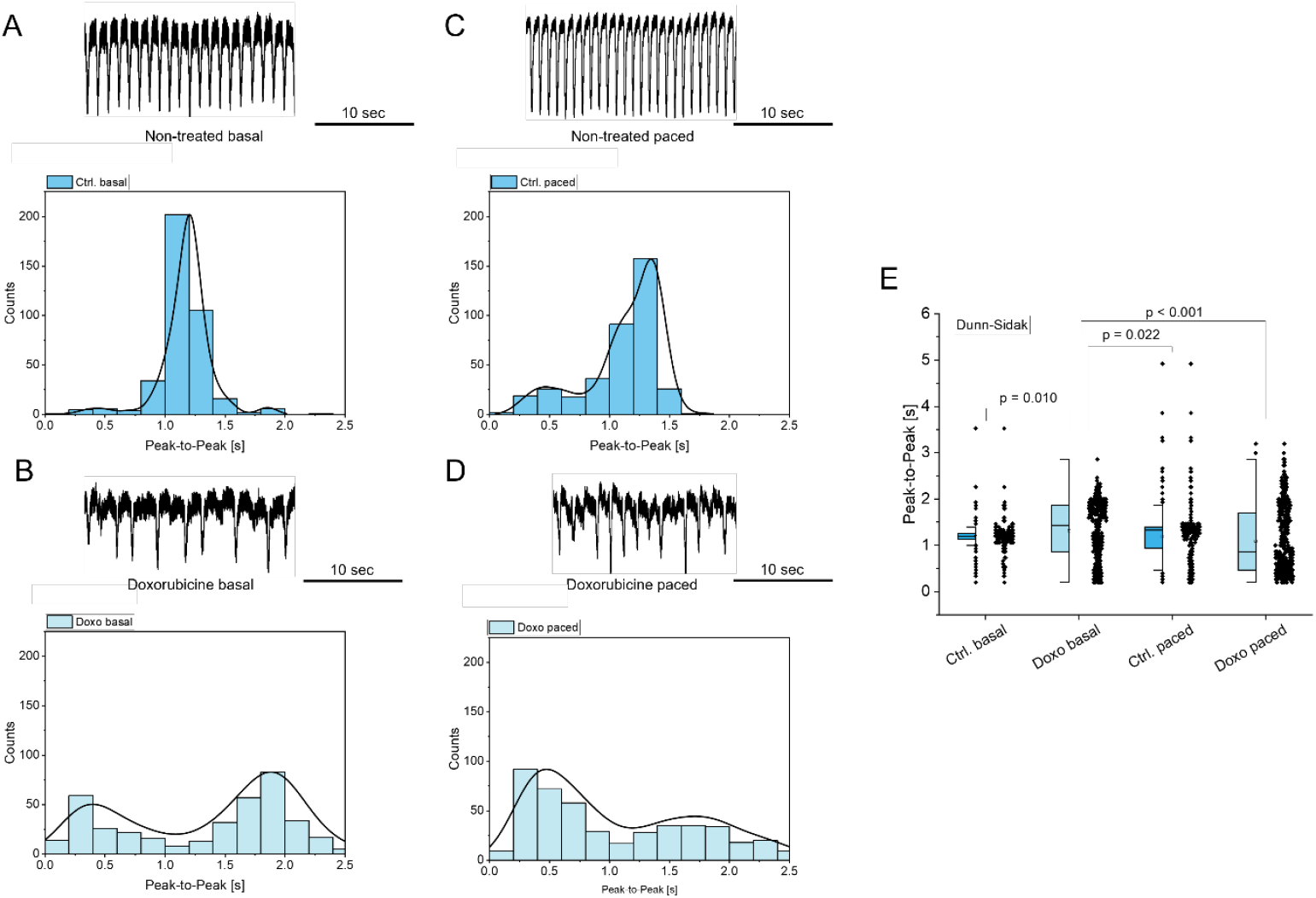
Doxorubicin-induced senescence induces contractile changes in hiPSC-derived ventricular CM. All measurements were done with the CardioExcyte 96 device (Nanion Technologies GmbH). **(A)** An example trace of the contraction of non-treated hiPSC-CM is shown on the top. The histogram and the plotted Kernel-Smooth-Fit shows a clear peak at 1.25 seconds peak-to-peak time. **(B)** Example trace of contractility of doxorubicin-treated hiPSC-CM. The histogram and the plotted Kernel-Smooth-Fit show two peaks at 0.25 seconds and 1.75 seconds peak-to-peak time. **(C)** Non-treated hiPSC-derived cardiomyocytes at 3 Hz pacing; an example trace is shown on the top. The histogram and the plotted Kernel-Smooth-Fit shows a clear peak at 1.3 seconds peak-to-peak time. Furthermore, the fit is slightly shifted to the left compared to **(A). (D)** Doxorubicin-treated hiPSC-CM at 3 Hz pacing. An example trace is shown on the top. The histogram and the plotted Kernel-Smooth-Fit show two peaks at 0.40 seconds and 1.75 seconds peak-to-peak time. **(E)**. Statistics: Dunn-Sidak, the vertical lines in the box blots represent the median values, the error bars denote the standard deviation (SD), the boxes represent the 25^th^ to 75^th^ percentile. n=2.

## 1. Discussion

Mitochondrial dysfunction and reduced respiratory capacity are well-known hallmarks of aging in different cell models and CM [52, 53]. We recently presented a CM senescence model derived from induced pluripotent stem cells and showed that this model behaves similarly to a senescence cell model of rat NVCM [54, 55]. We found reduced respiratory function as well as altered ultrastructure with an increase in truncated cristae [15]. With the present study, we provide a deeper analysis how cristae re-structuring and ATP synthase organization are related, compromising mitochondrial energy metabolism and eventually leading to irregular beating activity of senescent CM (Figure 6: Graphical abstract). First, we observed reduced mitochondrial ATP synthesis in our senescent hiPSC-CM model. Senescent CM tend to shift to increased glucose usage as their preferred fuel und thus exhibit a higher rate of glycolysis [13]. This has been attributed to age-related hyperglycemia, insulin resistance and type II diabetes, which are, together with aging, principal risk factors for the development of CVDs [56]. Although senescent CM had a higher proportion of glycolysis, we found decreased overall ATP production indicating that glycolysis could not compensate for the decrease in mitochondrial ATP production. While ATP production was decreased, ΔΨ_m_ was not changed in the senescent hiPSC-CM.

**Figure 6:**
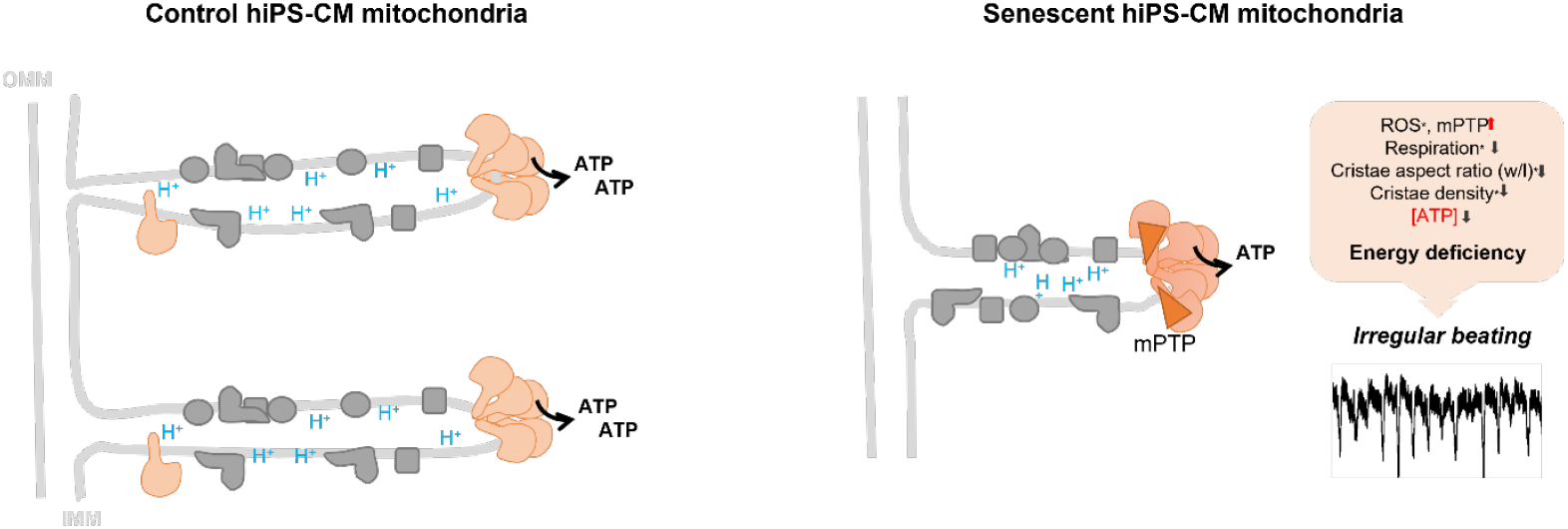
Graphical abstract.

Recently, it has been reported that aging is associated with a loss of ATP dimers [13, 40], which represent ATP synthase in the ATP synthesizing mode, while dissociation of dimeric and oligomeric ATP synthase is associated with reverse function [19]. However, we observed no changes in the ratio of monomeric, dimeric, and oligomeric ATP synthase in senescent NVCMs. Since neither the number nor the organization of ATP synthase complexes in monomers, dimers and oligomers was altered in our senescent models (hiPSC-CM and NVCM), we looked for other functions that could impact energy efficiency. ATP synthase is a reverse enzyme that can maintain the ΔΨ_m_ by pumping protons into the intra-cristae space, which results in acidification at subunit e as we recently showed [21]. To show this, a pH-sensor was genetically fused to the subunit e of ATP synthase, however, this assay was beyond the scope of the here presented study and at the moment we cannot answer the question whether ATP hydrolysis function of ATP synthase is responsible for the decreased ATP production. Third, ATP synthase is also thought to be involved in mPTP formation. Independ of which model is considered, the dimer model [57], the c-ring model [58] or the death-finger model [45, 57, 59], opening of the pore counteracts ATP synthesis. Indeed, mPTP opening was recently correlated with aging in mice heart [13] and our data suggest that mitochondria in our senescent CM model also exhibit increased mPTP, which would explain decreased ATP production.

Finally, we wondered whether the localization of ATP synthase, which is usually found at cristae rims, might be altered. This could affect the coupling with the ETC, the optimal utilization of the PMF and also mPTP opening. To determine the localization of ATP synthase in the IMM sub-compartments, cristae and inner boundary membrane, we performed tracking and localization microscopy (TALM). The course of trajectories allows to dissect, whether ATP synthase is mobile (monomers of ATP synthase show a higher mobility in the membrane than dimers and oligomers [42]) and reveals the microcompartments it occupies [47, 60]. Importantly, we observed significant changes in ATP synthase mobility in terms of step length and diffusion coefficient. Both parameters were significant lower in senescent conditions, suggesting either a reduced microcompartment or increased oligomerization. From the BN-data we have no indications for the latter. Also, others found rather a decrease than an increase in ATP synthase dimers and oligomers in aging conditions [13, 40]. Therefore, we propose that the altered ultrastructure of the IMM in senescent CM with a loss of cristae an increased number of truncated cristae and most important larger fenestration changes the cristae curvature significantly. Since ATP synthase has a preference for curved membrane parts, this would result in increased accumulation and likely oligomerization of ATP synthase in the curved parts [24, 61, 62], which would eventually result in decreased mobility. Such pronounced accumulation of ATP synthase, in addition to the formation of regular rows of dimers [63], may also favor the unstructured clustering of ATP synthase, which could be detrimental to ATP synthesis. Whether this is related to increased mPTP opening, is currently unclear and requires further investigation. For example, the increased ROS generation in senescent hiPS-CM [15] could be a trigger for mPTP opening [64]. Taken together, the impaired ATP synthesis in senescent CM provides a possible explanation for the observed irregularity of the contractile activity after stimulation in senescent CM (Figure 5). Under normal conditions, OXPHOS-linked ATP production may be sufficient, but under higher energy demands, this may not be possible due to a lack of extra capacity and increased mPTP. Lack of reserve capacity for ATP production and increased mPTP formation would contribute to age-related insufficiency of CM activity, particularly under energy-intensive conditions.

### Acknowledgements

We thank Paul Disse for helping growing and differentiating hiPS-CM.

## Authors’ contributions

SM, SP, GS, and VE and KBB designed the experiments, validated the data. SM, IMR, GB, FS, SP, KBB performed the experiments. SM, GS, VE and GB contributed to the methodology. KBB, GS and VE contributed to resources; SM and KBB wrote the manuscript; KBB, SM, GB and SP visualized the study; All authors supervised the data; KBB and VE contributed to project administration; KBB and VE acquired funding. All authors have read and agreed to the published version of the manuscript.

## Data availability statement

Raw data will be made available upon request.

