## supplementary figures for "Altered dynamics of ATP synthase cause functional irregularities in senescent hiPS-cardiomyocytes"

### Supplementary Material

#### Supplementary Figures

**Figure S 1: FCCP effect on $\Delta\Psi_m$: (**A) Typical time course showing uncoupling and changes of $\Delta\Psi_m$. (B) Quantification of FCCP induced $\Delta\Psi_m$ changes. n=1 biological replicate.

**Figure S 2: Respiratory chain complex activity assay.** hiPS-CM were permeabilized to give access to the substrates. Complex activity assay was performed via Seahorse extracellular flux analysis. OCR was recorded after subsequently adding substrate for CI (Mal/Glut), inhibiting CI with Rotenone, adding Succinate/G3P (CII substrate), Antimycin A (CIII inhibitor), and TMPD/Ascorbate (CIV substrate).

**Figure S 3: Expression of OXPHOS proteins is unaltered in senescent CM.** Expression of different complex subunits was determined via qPCR, normalized to GAPDH acted as housekeeping gene.

**Figure S 4: Fluorescence-tagging of ATP synthase in hiPSC-CM via HaloTag-fusion and labeling.** (A) Generation of a hiPS cell line expressing subunit γ-HaloTag. Immunostaining of BN-PAGE separated ATP synthase. (B) Successful differentiation into cardiomyocytes indicated by CM marker α-actinin. (C) Mitochondrial localization of subunit γ-HaloTag.

Figure S 1

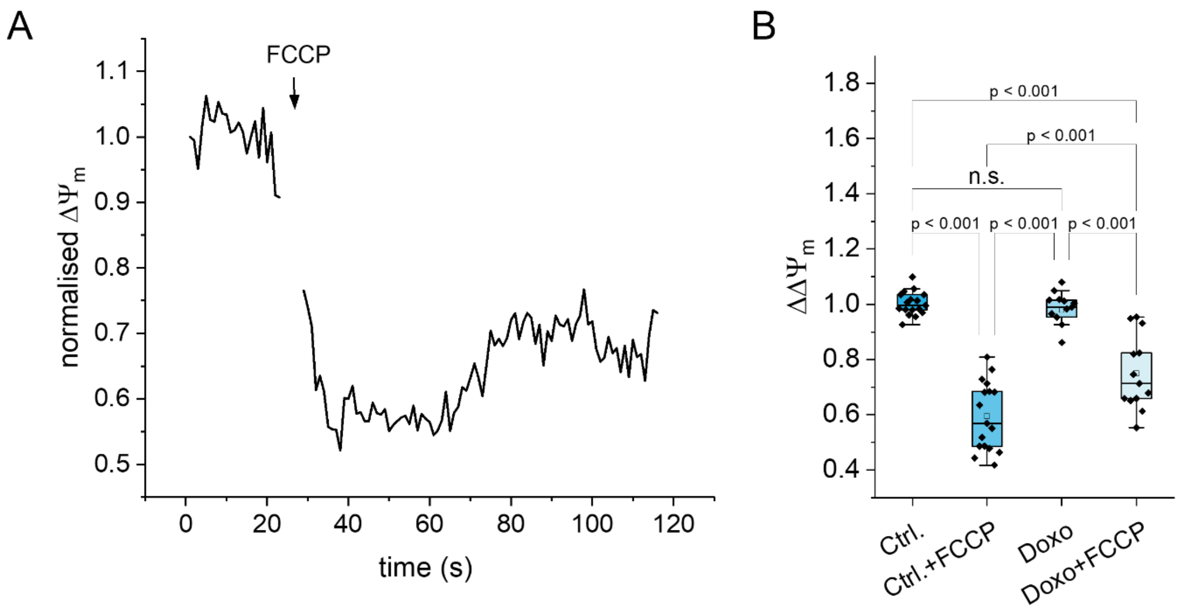

A
FCCP
normalised ΔΨm
1.1
1.0
0.9
0.8
0.7
0.6
0.5
0
20
40
60
80
100
120
time (s)
B
ΔΔΨm
1.8
1.6
1.4
1.2
1.0
0.8
0.6
0.4
p < 0.001
p < 0.001
n.s.
p < 0.001
p < 0.001
p < 0.001
Ctrl.
Ctrl.+FCCP
Doxo
Doxo+FCCP

Figure S 2

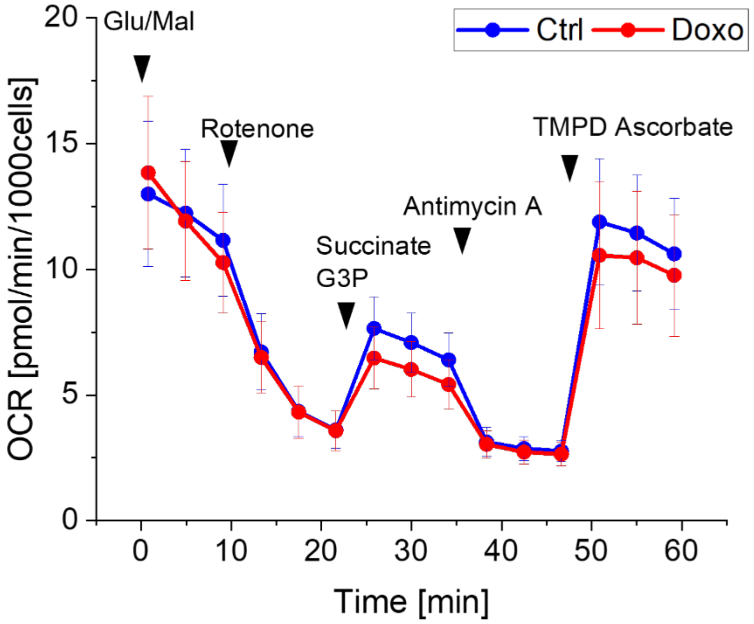

Glu/Mal
Rotenone
Succinate
G3P
Antimycin A
TMPD Ascorbate
Ctrl
Doxo
OCR [pmol/min/1000cells]
Time [min]
0
5
10
15
20
0
10
20
30
40
50
60

Figure S 3

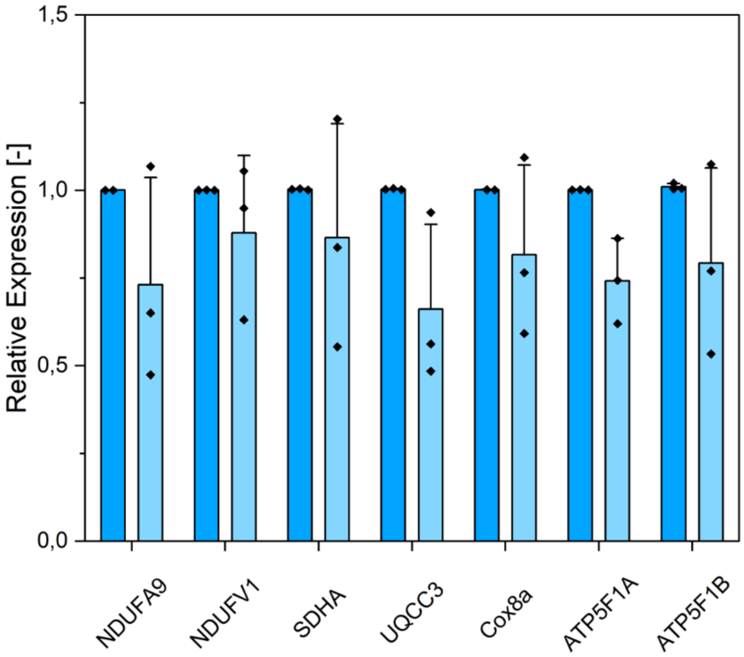

1,5
1,0
0,5
0,0
Relative Expression [-]
NDUFA9
NDUFV1
SDHA
UQCC3
Cox8a
ATP5F1A
ATP5F1B

### Figure S 4

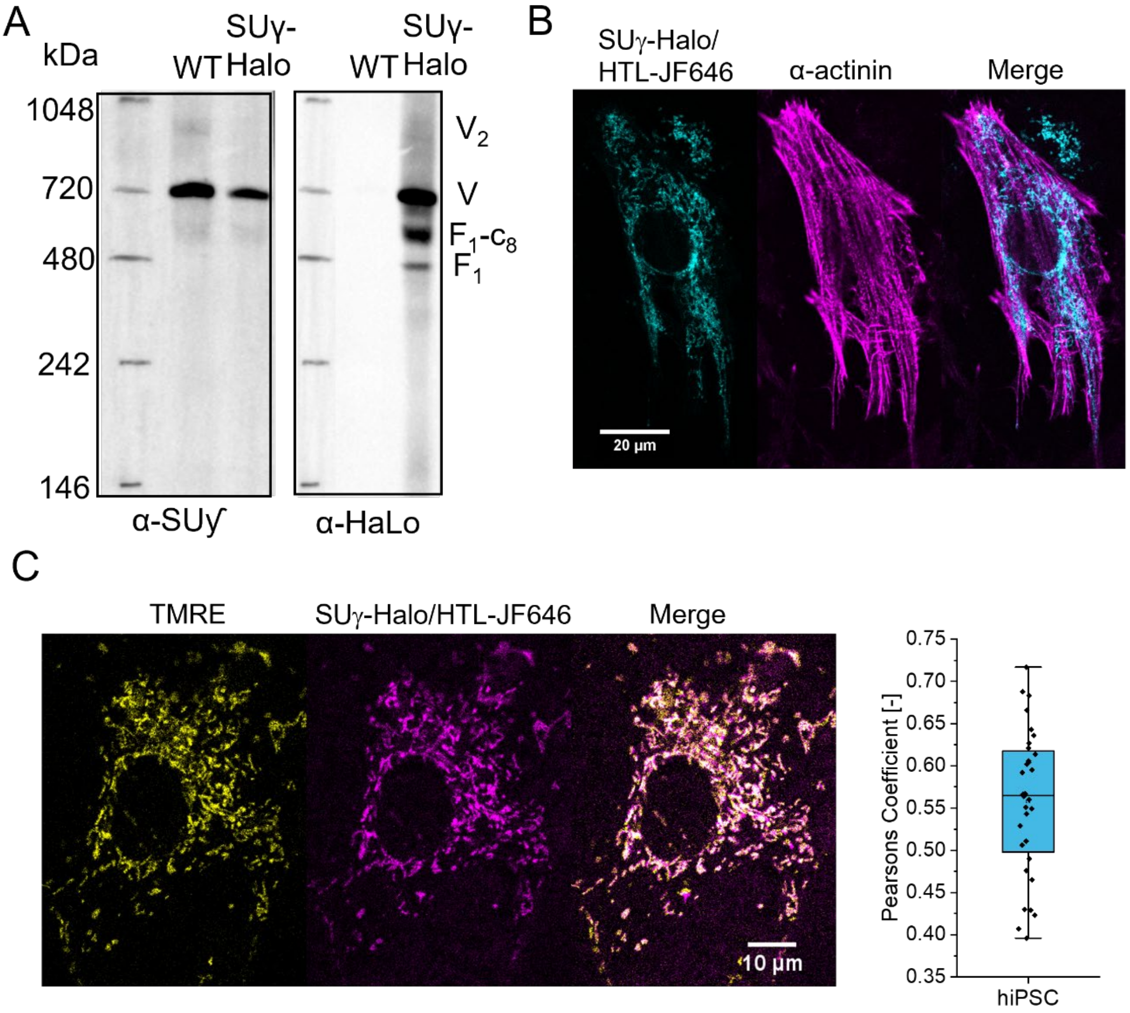
A
kDa
WT
SUγ-Halo
WT
SUγ-Halo
1048
720
480
242
146
V2
V
F1-c8
F1
α-SUγ
α-HaLo
B
SUγ-Halo/HTL-JF646
α-actinin
Merge
20 µm
C
TMRE
SUγ-Halo/HTL-JF646
Merge
10 µm
Pearsons Coefficient [-]
0.75
0.70
0.65
0.60
0.55
0.50
0.45
0.40
0.35
hiPSC
